# Towards Continuous Home Monitoring for Dementia: A Real-Time mmWave Radar Framework for Activity Classification and Tracking

**DOI:** 10.64898/2026.05.05.722929

**Authors:** Ziwei Chen, Charalambos Hadjipanayi, Maowen Yin, Alan Bannon, Timothy G. Constandinou

## Abstract

Millimeter-wave radar can quietly monitor health and behavior at home, which is vital for supporting people living with dementia. Most studies, however, remain limited to short-term testing in controlled spaces. Real-world deployment requires robust activity classification as a prerequisite: vital-sign and behavioral sensing require fundamentally different processing pipelines, and absent periods need to be reliably distinguished from stationary states. Bridging the critical gap between controlled laboratory demonstrations and continuous home monitoring, this paper introduces a self-adapting radar framework that extracts meaningful behavioral segments from massive, unconstrained real-world data. The system performs continuous real-time activity classification (stationary, walking, and absent) and target localization, selectively directing downstream processing to the most informative segments. It addresses key real-world deployment challenges including adaptive thresholding across subjects and environments, and walking detection under naturalistic activity conditions. Prior to integration with the Minder platform, the system was validated in a fully instrumented studio apartment against ground truth. Across 12 subjects, the system achieved an overall classification accuracy of 0.98, with F1 scores of 0.99 for absence and stationary states, and 0.95 for walking. Event-based evaluation yielded a per-subject walking sensitivity of 0.916*±* 0.058 and F1 score of 0.935 *±*0.030. Localization root mean square error during movement was 0.40 m. The results demonstrate reliable performance suitable for transitioning to long-term real-world home deployment.

## I. Introduction

**A**BOUT 85% of people living with dementia (PLWD) prefer to remain in their own homes for as long as possible [1], and home monitoring directly supports this preference.

Two-thirds of general practitioners (GPs) would prescribe assistive technologies to patients with early-stage dementia [2], and two-thirds of carers believe these technologies improve the quality of life of PLWD [3]. Independent living support enhances autonomy, improves safety with minimal restriction, boosts confidence, reduces caregiver burden, and preserves dignity.

Millimeter-wave (MmWave) radar offers a privacy-preserving and unobtrusive solution for continuous home monitoring of PLWD. As a platform technology, it enables both vital sign monitoring and behavioral sensing, providing advance warning of disease progression, including vital sign changes, frailty, gait instability, and adverse events such as falls. Most importantly, radar operates autonomously 24/7 without the privacy concerns of cameras or microphones. It also overcomes the key limitations of wearable sensors, as sustained contact causes discomfort and may alter the behaviors being monitored. Elderly users often struggle with device maintenance and recharging, further disrupting continuous recording [4] [5].

However, if radar research continues to rely on short-term studies in controlled environments, the real-world advantages of this technology remain unrealized. Despite significant progress, the majority of radar studies rely on short-duration experiments featuring predefined and scripted activities [6]. For example, radar-based activity recognition systems have achieved high classification accuracy but are typically trained and evaluated on scripted activity sequences under controlled conditions [7] [8]. Hsu et al. [9] demonstrated that RF signals can support a range of home sensing tasks including gait analysis, sleep monitoring, and user identification, but each was developed and validated as a separate system. A primary advantage of radar monitoring, however, is the ability to track disease progression continuously over months or years, rather than relying on sporadic hospital visits.

Real-world radar data collection faces several practical hurdles. Reliable and privacy-preserving ground truth is difficult to obtain in 24/7 home scenarios, and studies involving patients across diverse home environments require substantial funding. Furthermore, the vast raw data volume generated by continuous monitoring presents logistical challenges in storage and post-processing, making long-term deployment difficult to scale [10]. Consequently, researchers often resort to condensed activity sequences over short periods to maximize event density for machine learning (ML) training and validation, raising concerns about the generalizability of those models in real-world home deployments.

The UK Dementia Research Institute (UKDRI) aims to discover, develop and deliver solutions to maintain brain health in an aging society. Its Care Research and Technology center is developing the Minder platform, which currently uses home sensors across 100 private residences to gather continuous environmental, physiological, and activity data in unprecedented detail to support dementia care [11]. Integrating radar technology into Minder presents a valuable opportunity to access real patient data in genuine home environments, but it also comes with implementation challenges. Under General Data Protection Regulation (GDPR), healthcare-related data are highly sensitive, and thus privacy-intrusive devices such as cameras are excluded from the implementations. Moreover, raw data cannot be saved or uploaded directly at this stage due to its volume, and only small snippets could realistically be handled, so a practical method is needed to decide what data to retain and compress. This also means that conventional approach of recording, uploading and manually labeling raw data is not feasible. Therefore, efficient on-site strategies are required to handle and extract meaningful information from large data volumes in real time over continuous monitoring periods.

Beyond the challenge of data collection, two further radar processing challenges arise. First, vital sign monitoring and behavioral sensing require fundamentally different processing pipelines: vital sign measurement results are only valid when the subject is stationary, and gait analysis is only meaningful during walking [10] [12]. Second, not all 24/7 data are informative. Absent periods are less informative and energy consuming, yet common in 24/7 deployment. They share similar radar signatures with stationary periods, particularly for distant or lying down subjects, where both yield empty range-Doppler maps and weak chest-wall motion [13] [14]. Although target tracking algorithms have been widely explored in radar research, most function as a motion tracker, and revert to an absent state when the target stays stationary [15]. While in those applications stationary periods are not the main concern, this is inadequate for 24/7 home monitoring: where stationary periods, such as watching TV, reading, or napping in various indoor locations, are common, highly informative, and critical for real-life applications. False presence also introduces significant waste of storing and processing effort. In our application, the radar generates approximately 80 GB of compressed raw data per 10-hour period. Hence, a robust target activity classifier is essential to correctly distinguish absent from stationary states and ensure processing effort is directed where it is most useful.

This work proposes an adaptive radar framework that elevates a basic proof-of-concept live demonstration into a fully deployable system for continuous activity classification and target localization in actual homes. To build this architecture, the earlier ultra wideband (UWB) system is replaced with a 60 GHz frequency-modulated continuous-wave (FMCW) radar, which enables 2D spatial tracking, widespread availability, and ease of integration into compact embedded systems [16]. The algorithm selectively directs processing resources to the most informative segments: stationary periods for vital sign monitoring, walking periods for gait analysis and subject identification, and movement trajectory for tracking daily routines (Fig. 1). It also addresses key challenges in real-world deployment: robust thresholding that works for different subjects in various home environments, reducing false absence for stationary targets, and reliable walking detection during daily activity events.

**Fig. 1.**
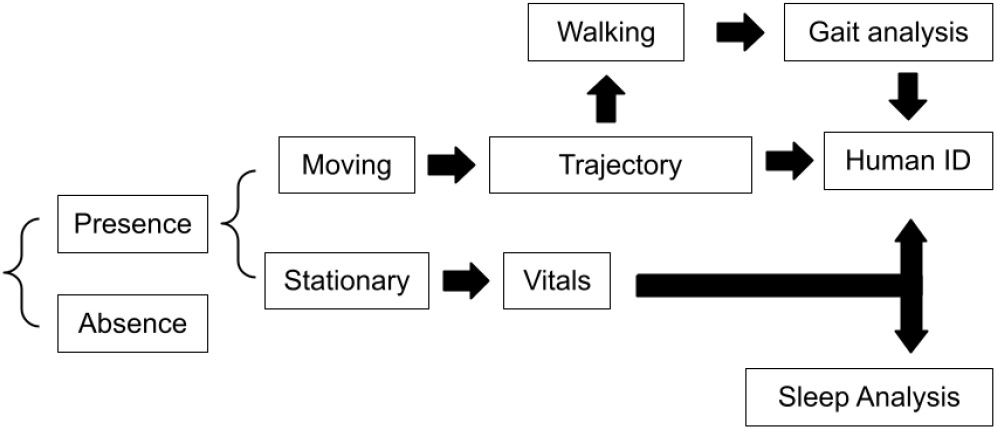
System overview.

The main contributions of this work are as follows:

- A unified real-time radar framework that classifies target activity (stationary, walking, and absent) and selectively directs downstream processing accordingly, enabling efficient continuous monitoring.
- A parallel preprocessing design with separate paths for motion-based tracking and breathing-based absence detection, with adaptive clutter removal and zero-Doppler range profile extraction respectively.
- A walking detection algorithm designed for naturalistic activity conditions, including variable-pace movement and unprompted directional changes.
- A 12-subject validation study conducted in a fully instrumented studio apartment with SensFloor and camera ground truth under naturalistic activity conditions.

## II. Methods

The 60 GHz Infineon FMCW radar (BGT60TR13C) development board connected to a laptop via USB is used, and processing is performed in Python. The radar is configured (Table I) to support a maximum detection range of 9.6 m, a range resolution of 0.05 m, and a maximum unambiguous Doppler velocity of 6.25 m/s. Two receivers are activated instead of all three to reduce data volume, and the azimuth plane is sufficient for tracking requirements.

**TABLE I.** FMCW Radar Configuration Parameters.

| Parameter Name | Value |
| --- | --- |
| Sampling rate | 2 MHz |
| Number of chirps per frame | 64 |
| Number of samples per chirp | 360 |
| Start frequency | 58 GHz |
| End frequency | 61 GHz |
| Frequency bandwidth | 3 GHz |
| Chirp duration time | 188.96 $\mu$ s |
| Frame duration time | 12.8 ms |
| Activated receivers | RX2 & RX3 |

The FMCW radar transmits frequency-varying chirps. Each reflected chirp is mixed with the transmitted signal, producing a beat signal whose frequency is proportional to target range. A first fast Fourier transform (FFT) along fast time hence yields the range profile. Across successive chirps, a moving target induces a progressive phase shift at each range bin, whose rate corresponds to its radial velocity. A second FFT along slow time thus produces the range-Doppler map (Fig. 2a). Three consecutive frames are concatenated to increase the slow-time aperture, yielding a velocity resolution of 0.065 m/s at an effective processing rate of 26 Hz.

**Fig. 2.**
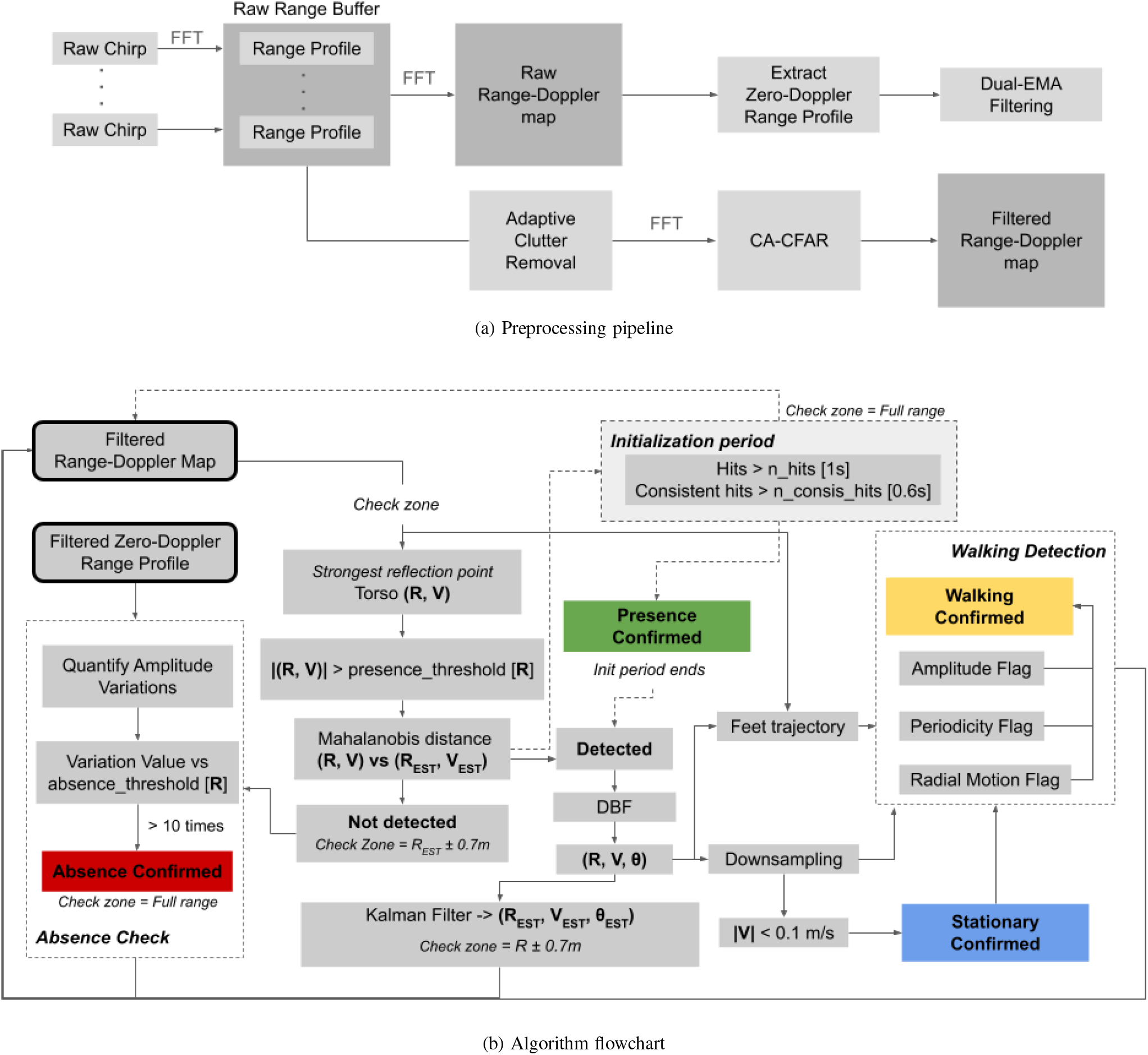
Flowchart for the mmWave radar framework.

As illustrated in Fig. 2a, pre-processing follows two parallel paths. The main tracker path applies the adaptive clutter removal method in [17] before the Doppler FFT, suppressing static clutter in the range domain to prevent spectral leakage into adjacent Doppler bins, followed by cell-averaging constant false alarm rate (CA-CFAR) detection. The absence check path instead uses the raw range-Doppler map (without zero-padding), because breathing energy is attenuated by clutter removal. The Doppler frequency resolution is Δ*f* =1*/*(*NT*_*c*_) ≈ 27 Hz, where *N* = 192 is the total number of chirps across the concatenated frames and *T*_*c*_ = 188.96 µs is the chirp repetition interval. Since this far exceeds typical breathing rates, the respiratory energy concentrates in the zero-Doppler bin. The zero-Doppler range profile is therefore used for absence check.

The radar framework consists of three main modules: (a) the main tracker, (b) the absence check, and (c) the walking detection (Fig 2b). The main tracker activates when a candidate torso point consistently exceeds the amplitude threshold. During active presence, it continuously records the subject’s location and extracting foot trajectory for walking detection. Periods with low torso speed are classified as stationary. If the number of valid targets falls below a threshold over recent frames, the system enters the absence check phase, where a drop in breathing band energy below threshold deactivates the tracker and returns the system to initialization.

### 1) Tracker

target tracking relies on range-dependent thresholding and Kalman filtering to construct daily activity trajectory. Target tracking is performed primarily in the range-Doppler domain, which provides substantially higher resolution than the angular resolution due to the single transmit antenna. Specifically, the strongest reflection point on the clutter-filtered range-Doppler map is selected as a candidate torso reflection. This candidate is confirmed as a valid target detection when it satisfies two criteria: (a) its amplitude is sufficiently large, and (b) it remains temporally stable across consecutive frames.

To address the first criterion, a range-dependent presence threshold is established. This is determined via a hardware-level calibration, ensuring that it is generalizable across different subjects and home environments. According to the monostatic radar equation [18], the received power of a radar return decreases inversely with the fourth power of the target range (1*/R*^4^). When expressed within the radar’s configured 9.6 m field-of-view (FoV), this relationship approximates a linear decay. Consequently, linear regression was applied to the reflection power versus range using data from three walking subjects across four room environments (healthy subjects aged 28-30, one female and two males, three home apartments and one corridor, excluding the validation room). This calibration yielded a consistent slope of approximately −5.6, to which an empirical offset of −79 dB was also added to suppress low-level noise (Fig. 3a). While this does not account for all variations in the absolute reflection amplitudes, it provides sufficient tracking accuracy by effectively normalizing amplitude fluctuations attributable to range. The threshold is calibration-free, as after clutter removal, the amplitude profile of a walking target follows the radar range equation, shaped by antenna gain and path loss.

**Fig. 3.**
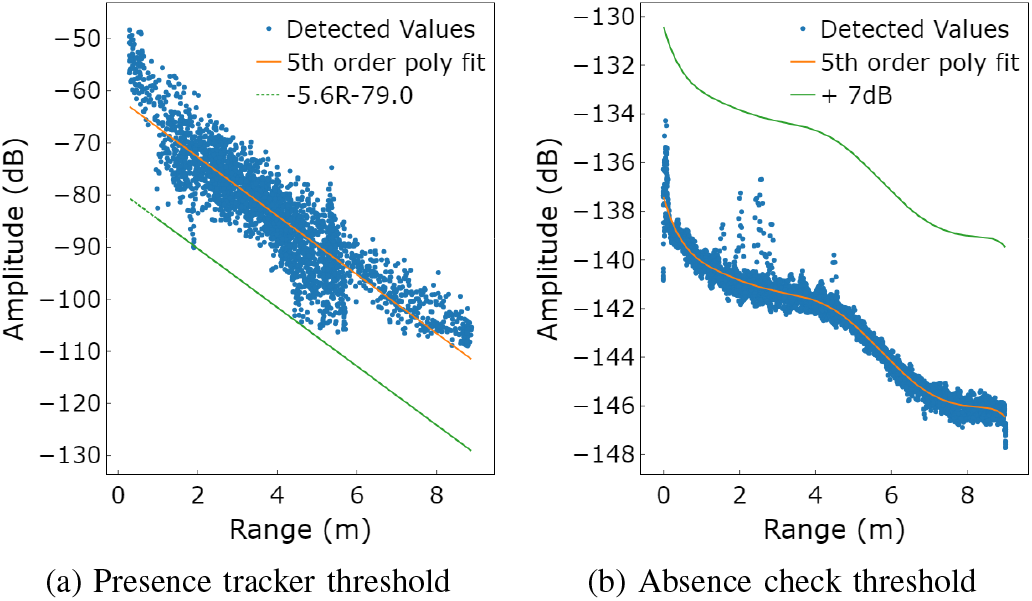
Calibration-free range-dependent detection thresholds: (a) presence tracker, (b) absence check.

Temporal stability is evaluated by computing the Mahalanobis distance between the newly detected target candidate and the predicted target state derived from the Kalman filter. By incorporating the state covariance matrix, the Mahalanobis distance effectively accounts for correlations between the state variables and naturally scales heterogeneous units, such as range and Doppler velocity [19]. Therefore, it is preferred over the Euclidean distance for data association and tracking within the range-Doppler domain.

Finally, digital beamforming (DBF) is applied across the receivers to produce an angular energy profile at the estimated target range. The peak azimuth is extracted and combined with range to obtain a 2D position estimate.

### 2) Absence Check and Stationary Target

as discussed in the introduction, continuous 24/7 monitoring necessitates a robust and generalizable method to distinguish between a stationary subject and true absence. The solution relies on isolating the minor chest-wall movements caused by human respiration. To achieve this, a dual exponential moving average (EMA) filter is applied to the zero-Doppler target range bin to isolate breathing energy [20]. This method is computationally lightweight, leverages historical data, and enables robust tracking of stationary subjects without triggering false absences.

In this work, EMA smoothing factors (weights) of 0.5 and 0.95 are adopted, instead of the 0.9 and 0.99 values proposed in [20]. While the latter values preserve more breathing energy, they exhibit significantly slower adaptation speeds. This is acceptable when extracting respiration from a known and perpetually stationary target. However, continuous in-home monitoring demands faster responsiveness to accommodate frequent transitions between moving, stationary, and absent states. The selected weight pair provides an optimal trade-off, ensuring faster adaptation speed while preserving sufficient breathing energy for reliable detection.

Consequently, a range-dependent absence threshold is estimated from filtered empty-room data (excluding the validation environment). It is calibration-free as dual-EMA filtering also suppresses reflections from static objects such as walls and furniture. The residual signal in an empty room is therefore dominated by the receiver noise floor. Both absence and presence thresholds are hardware-dependent and are expected to generalize across typical indoor environments without per-site recalibration, provided the same radar hardware is used.

The filtered breathing energy is quantified via maximum ratio combining (MRC) across range bins within a 2-second temporal window centered on the target range bin. The resulting value is compared against the absence threshold to determine target presence (Fig. 4).

**Fig. 4.**
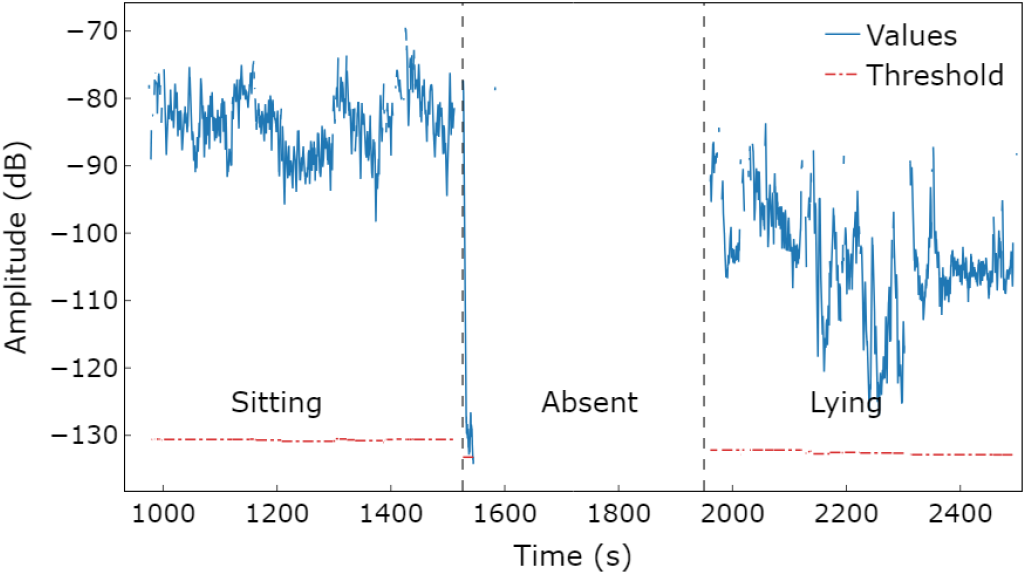
Breathing band energy during sitting, absent, and lying.

The initialization period of the presence-tracking state and the absence-check procedure introduce inherent temporal delays, but these durations are deterministic and can be precisely calculated and compensated for.

### 3) Walking Detection

walking periods benefit gait analysis and contain unique subject-specific information. Valid walking events are characterized by distinct radial motion, excluding in-place movements, and exhibit strong and periodic foot trajectory, which is also crucial for robust gait analysis [17]. A valid walking period is therefore determined by evaluating three sequential criteria: (a) amplitude, (b) periodicity, and (c) radial motion (Algorithm 1).

#### Algorithm 1 Walking Detection

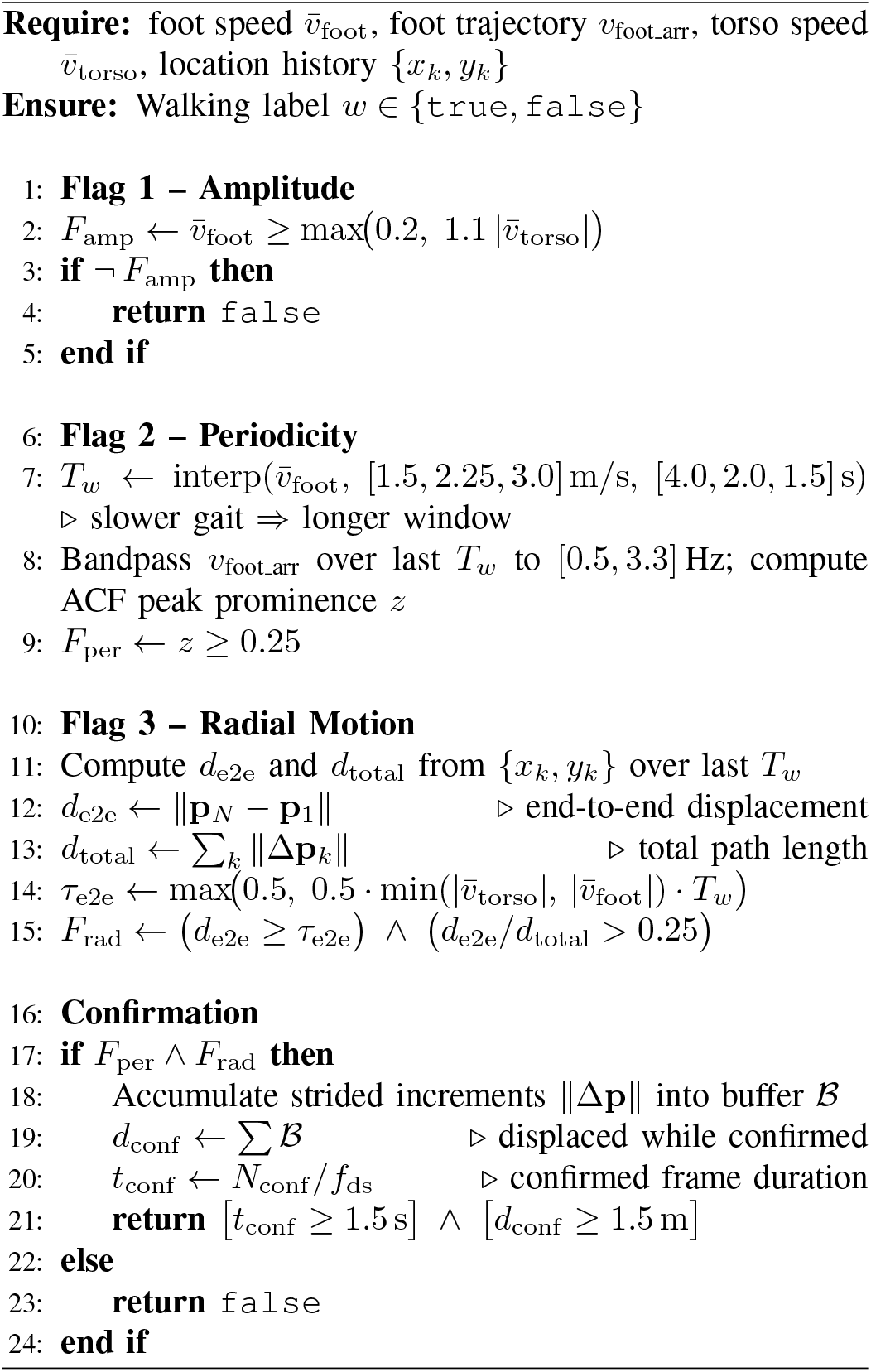

When walking, the fastest point on the range-Doppler map corresponds to the foot motion, and a continuous foot trajectory is extracted by isolating this peak in each frame [17]. To satisfy the amplitude flag, the instantaneous foot speed must exceed the torso speed by a factor of 1.1, with a minimum floor of 0.2 m/s to reject noise during near-stationary periods. To evaluate the periodicity flag, the windowed foot trajectory is band-pass filtered within the human cadence range (0.5-3.3 steps/s, spanning slow pathological to brisk healthy gait [21]), and the prominence value (*z* ≥ 0.25) of the resulting autocorrelation function (ACF) peak is then used to confirm cyclic gait patterns.

Window lengths range from 1.5 s at brisk speeds to 4.0 s at slow pathological gait. For the elderly group, slow walking speeds often dominate; however, a fixed long window would introduce excessive latency and risk missing the brief, intermittent walking bouts typical of daily in-home activity. Radial motion is confirmed by requiring sufficient end-to-end spatial displacement consistent with the total integrated distance traveled, with an end-to-end to total path length ratio (d_*e*2*e*_*/d*_*total*_) of at least 0.25, ensuring directional progression rather than stationary fidgeting. Finally, an overarching threshold (1.5 s duration, 1.5 m displacement) discards excessively brief walking bouts (short in duration or total distance moved), since these data provide insufficient information for reliable gait extraction and inflate the false positive rate. Example foot trajectories for structured and unstructured daily walking are shown in Fig. 5.

**Fig. 5.**
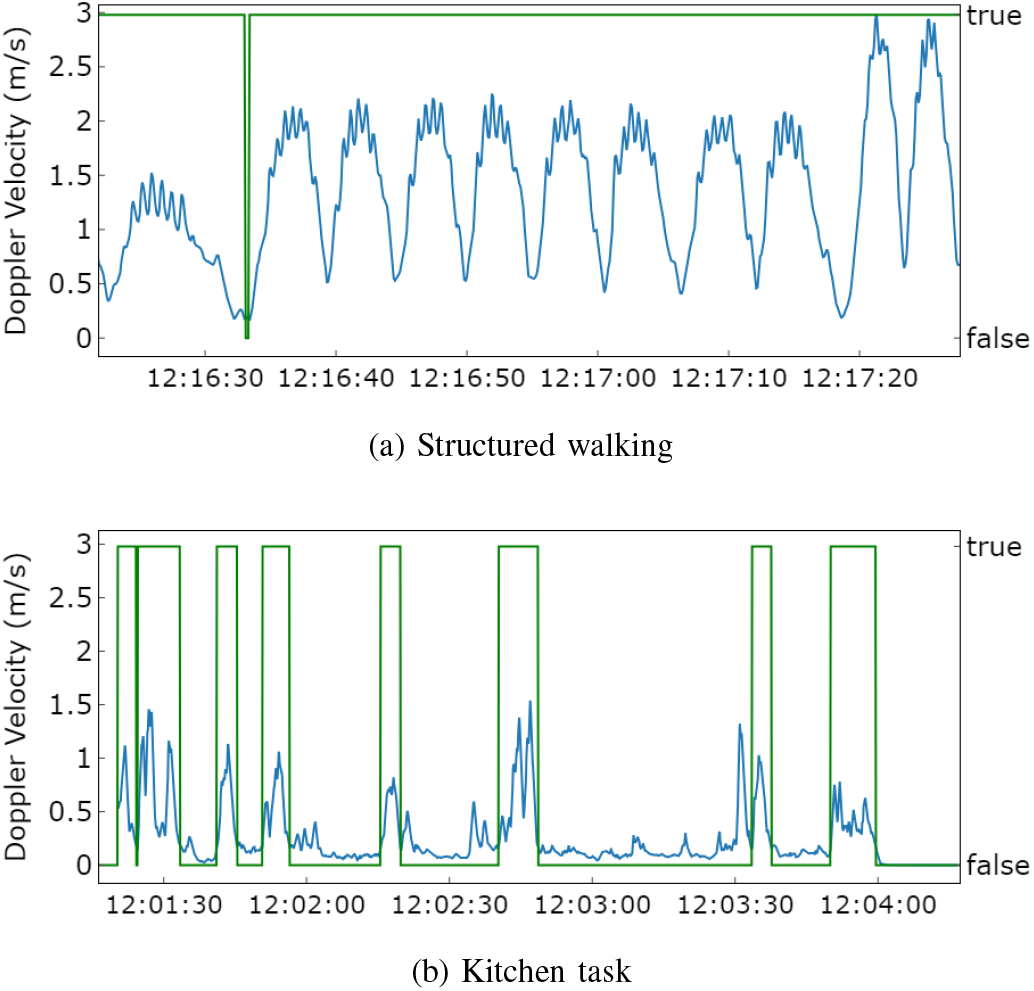
Foot trajectory (blue) and walking detection results (green): (a) structured walking, (b) short walking bouts during daily activities (kitchen task in Fig. 7).

## III. Validation

Prior to Minder deployment, where non-privacy-preserving measurements cannot be deployed in patient homes, a validation study was conducted in a fully instrumented studio apartment equipped with cameras (Microsoft Azure Kinect DK) and floor sensors (SensFloor Future-Shape GmbH) [22], as shown in Fig 6. Each SensFloor module measures 0.5 × 0.5 m and contains eight triangular proximity fields, reporting wirelessly at 10 Hz. All system parameters and thresholds were set using data from outside the validation environment, demonstrating generalizability to unseen subjects and settings. This work was approved by the Imperial College’s Science, Engineering and Technology Research Ethics Committee (SETREC number: 7365801).

**Fig. 6.**
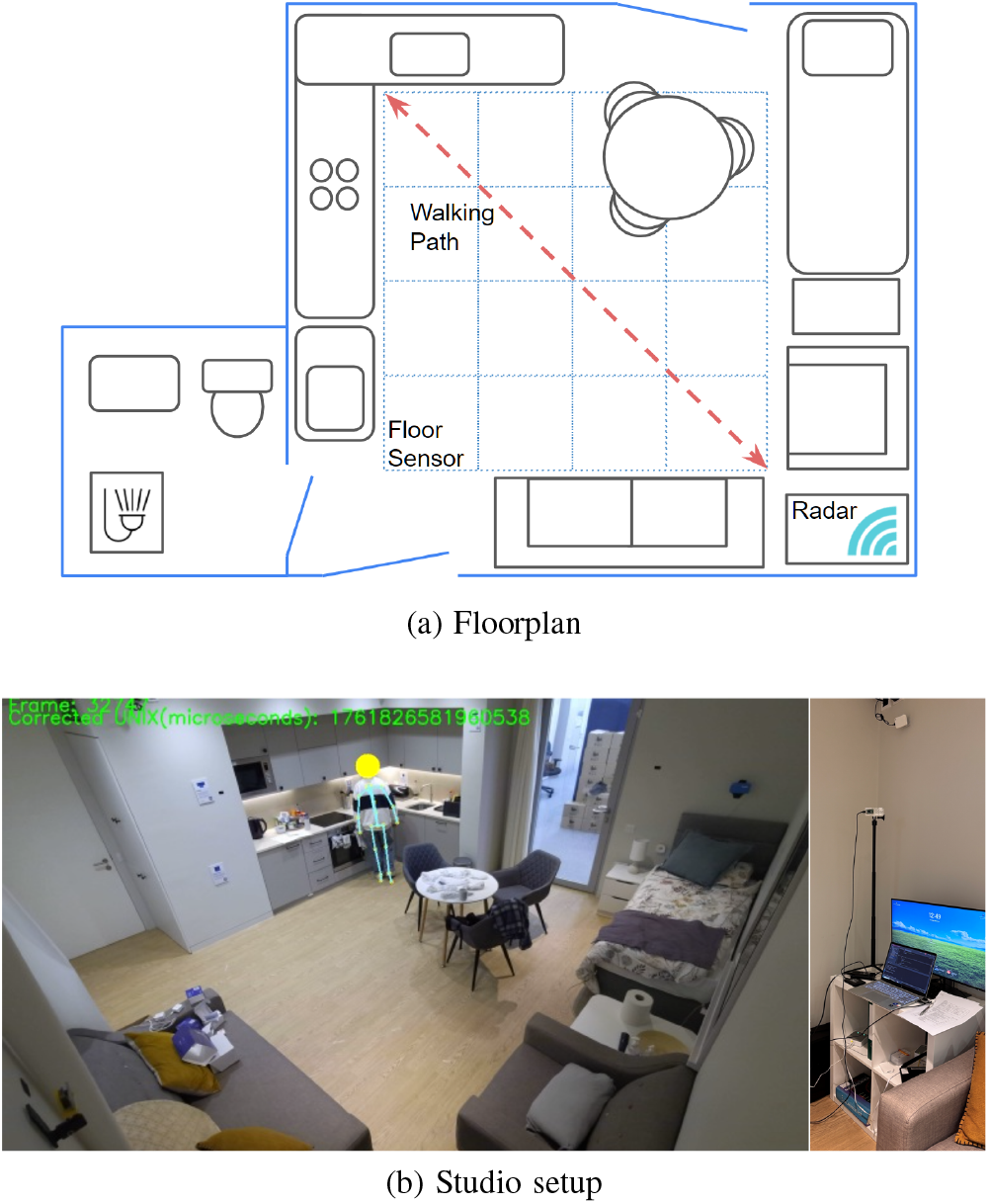
Experimental setup: (a) floorplan with radar placement (height=1.7 m) and walking path, (b) studio apartment photograph.

Twelve healthy participants (6 male, 6 female; aged 23-32, mean 27) completed a randomized protocol of nine activities across four activity sessions (Fig. 7). Activity order, sofa, chair, and standing positions were pre-assigned and counter-balanced to ensure spatial coverage of the monitoring area. The activities were designed to cover boundaries between states, including long-duration stationary periods with varied postures and locations, absence intervals, and free exploration tasks.

**Fig. 7.**
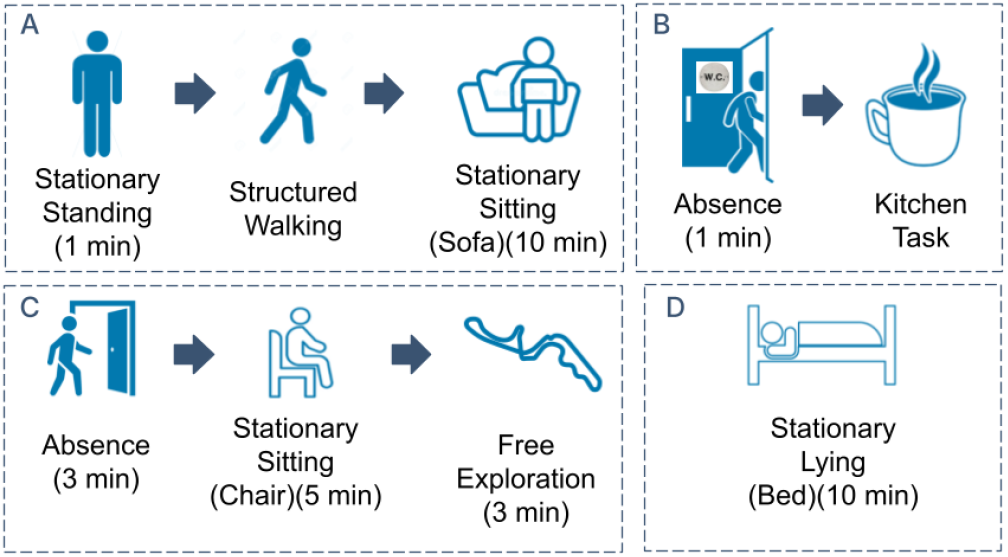
Experimental protocol activity sessions.

Session A established a baseline with a 1-minute standing phase, followed by structured walking at slow, normal, and fast speeds (participants carried a full cup of water during the slow phase to naturally reduce pace), and a prolonged sitting period on one of the sofas. Session B comprised a short absence interval (toilet visit), followed by a kitchen task (preparing a hot beverage) involving walking and upper-body movements. Session C included a longer absence phase (leaving through the main door), re-entry and relaxation on a chair to assess re-acquisition and stationary detection, concluding with a free exploration segment in which participants searched for a hidden object. Session D involved a lying phase on the bed, with participants instructed to transition among supine, right-side, and left-side postures every three minutes.

## IV. Results

The performance of the system is evaluated in terms of continuous activity classification against ground truth and target localization during various movement phases.

### 1) Classification

classification results were obtained from manually labeled ground truth derived from camera data. An example classifier output is shown in Fig. 8. While stationary states were marked per frame (10 Hz), only stationary periods lasting longer than 25 seconds were selected for evaluation and confirmed against the camera data.

**Fig. 8.**
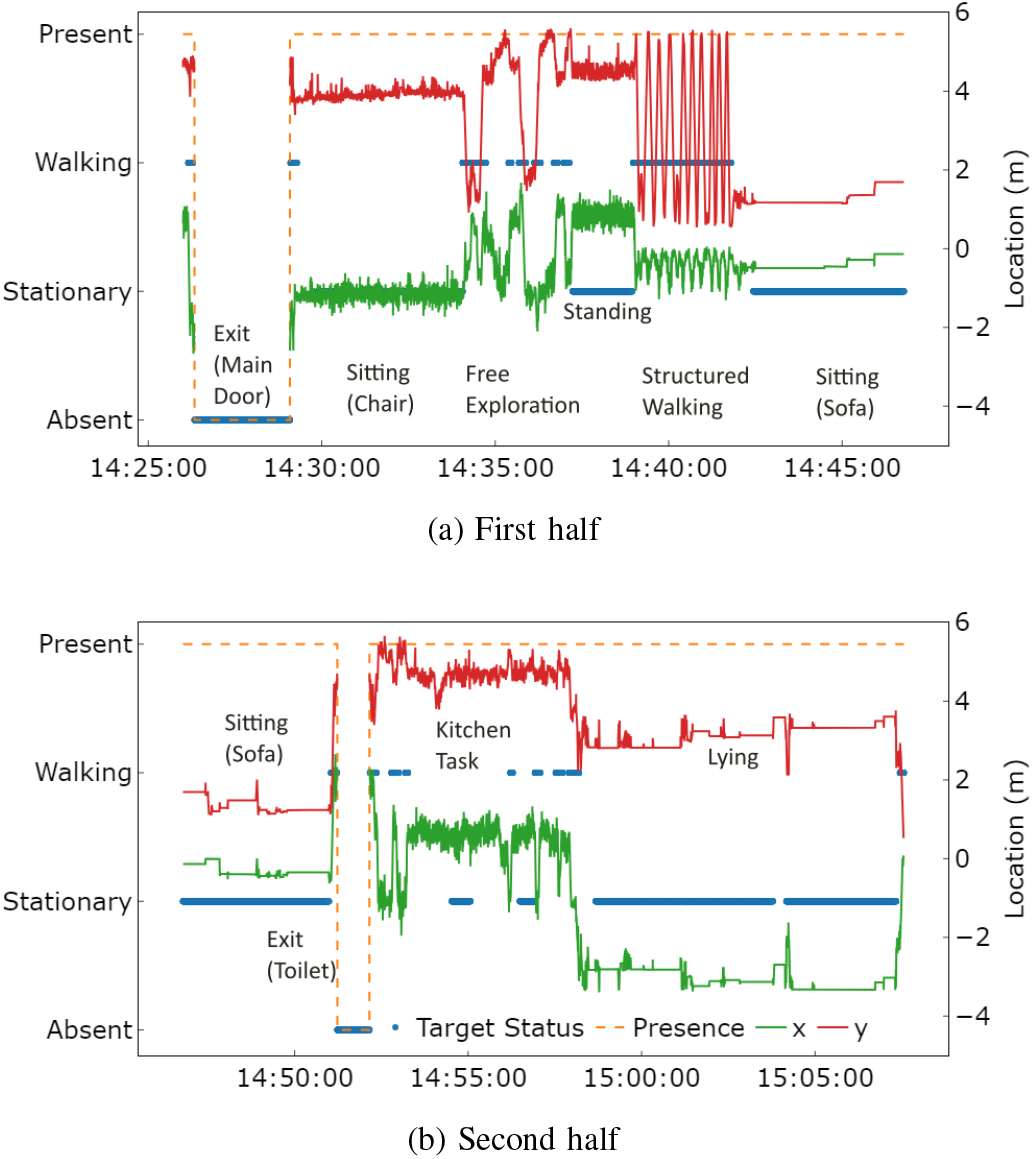
Classifier output showing target state and location across a continuous monitoring session (activity sequence CABD in Fig. 7): (a) first half and (b) second half of the session.

Table II presents the frame-level classification results evaluated at 10 Hz across all 12 subjects (N = 230,045 frames). The system achieved an overall accuracy of 98%, with F1 scores of 0.99, 0.99, and 0.95 for the absent, stationary, and walking classes respectively. The primary source of misclassification was within the walking class, where 2,818 frames were assigned to the other presence category and 16 to stationary.

**TABLE II.** Frame-based Confusion Matrix and Classification Metrics (Downsampled to 10 Hz)

| True State | Predicted State |  |  |  | Metrics |  |  |  |
| --- | --- | --- | --- | --- | --- | --- | --- | --- |
|  | Absent | Stationary | Walking | Other | Precision | Recall | F1 Score | Support |
| Absent | 27,667 | 0 | 157 | 316 | 1.00 | 0.98 | 0.99 | 28,140 |
| Stationary | 0 | 165,390 | 51 | 2,011 | 1.00 | 0.99 | 0.99 | 167,452 |
| Walking | 0 | 16 | 31,619 | 2,818 | 0.99 | 0.92 | 0.95 | 34,453 |
| Overall Accuracy |  |  |  |  |  |  | **0.98** | 230,045 |

Table III presents the event-based evaluation. At the aggregate level, walking detection achieved a sensitivity of 0.91, precision of 0.96, and F1 score of 0.93. The duration-weighted sensitivity (*S*_*d*_) was 0.92, where this metric is defined as the ratio of correctly detected walking time to the total ground-truth walking duration. Stationary and absent events were detected with high sensitivity and F1 scores of 1.00 and 0.98 respectively. Per-subject results (Table III Part B) confirm the consistency of these findings across individuals: walking sensitivity was 0.916± 0.058 and precision 0.959 ±0.042, while stationary and absent events showed near-perfect sensitivity with negligible variance (1.000 ± 0.000 for both).

**TABLE III.**
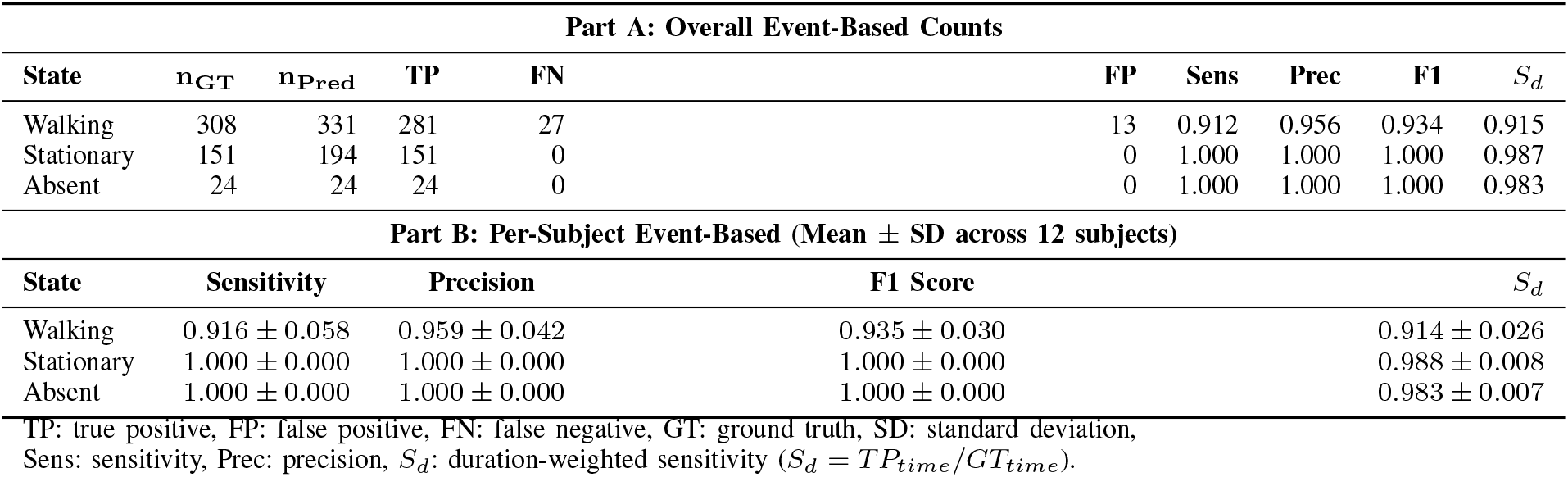
Event-Based Evaluation and Per-Subject Performance.

| Part A: Overall Event-Based Counts |  |  |  |  |  |  |  |  |  |
| --- | --- | --- | --- | --- | --- | --- | --- | --- | --- |
| State | n<sub>GT</sub> | n<sub>Pred</sub> | TP | FN | FP | Sens | Prec | F1 | *S<sub>d</sub>* |
| Walking | 308 | 331 | 281 | 27 | 13 | 0.912 | 0.956 | 0.934 | 0.915 |
| Stationary | 151 | 194 | 151 | 0 | 0 | 1.000 | 1.000 | 1.000 | 0.987 |
| Absent | 24 | 24 | 24 | 0 | 0 | 1.000 | 1.000 | 1.000 | 0.983 |
| Part B: Per-Subject Event-Based (Mean ± SD across 12 subjects) |  |  |  |  |  |  |  |  |  |
| State | Sensitivity |  | Precision |  | F1 Score |  | *S<sub>d</sub>* |  |  |
| Walking | 0.916 ± 0.058 |  | 0.959 ± 0.042 |  | 0.935 ± 0.030 |  | 0.914 ± 0.026 |  |  |
| Stationary | 1.000 ± 0.000 |  | 1.000 ± 0.000 |  | 1.000 ± 0.000 |  | 0.988 ± 0.008 |  |  |
| Absent | 1.000 ± 0.000 |  | 1.000 ± 0.000 |  | 1.000 ± 0.000 |  | 0.983 ± 0.007 |  |  |

TP: true positive, FP: false positive, FN: false negative, GT: ground truth, SD: standard deviation, Sens: sensitivity, Prec: precision, S<sub>d</sub>: duration-weighted sensitivity ( $S_d = TP_{time}/GT_{time}$ ).

### 2) Localization

the floor sensor generates no position output during stationary phases, therefore evaluation focused on three activity segments: kitchen task, structured walking, and free exploration. To correct accumulated time offsets between sensors, each segment was manually peak-aligned before evaluation. Fig. 9a and 9b show example trajectories and range-time profiles during structured walking.

**Fig. 9.**
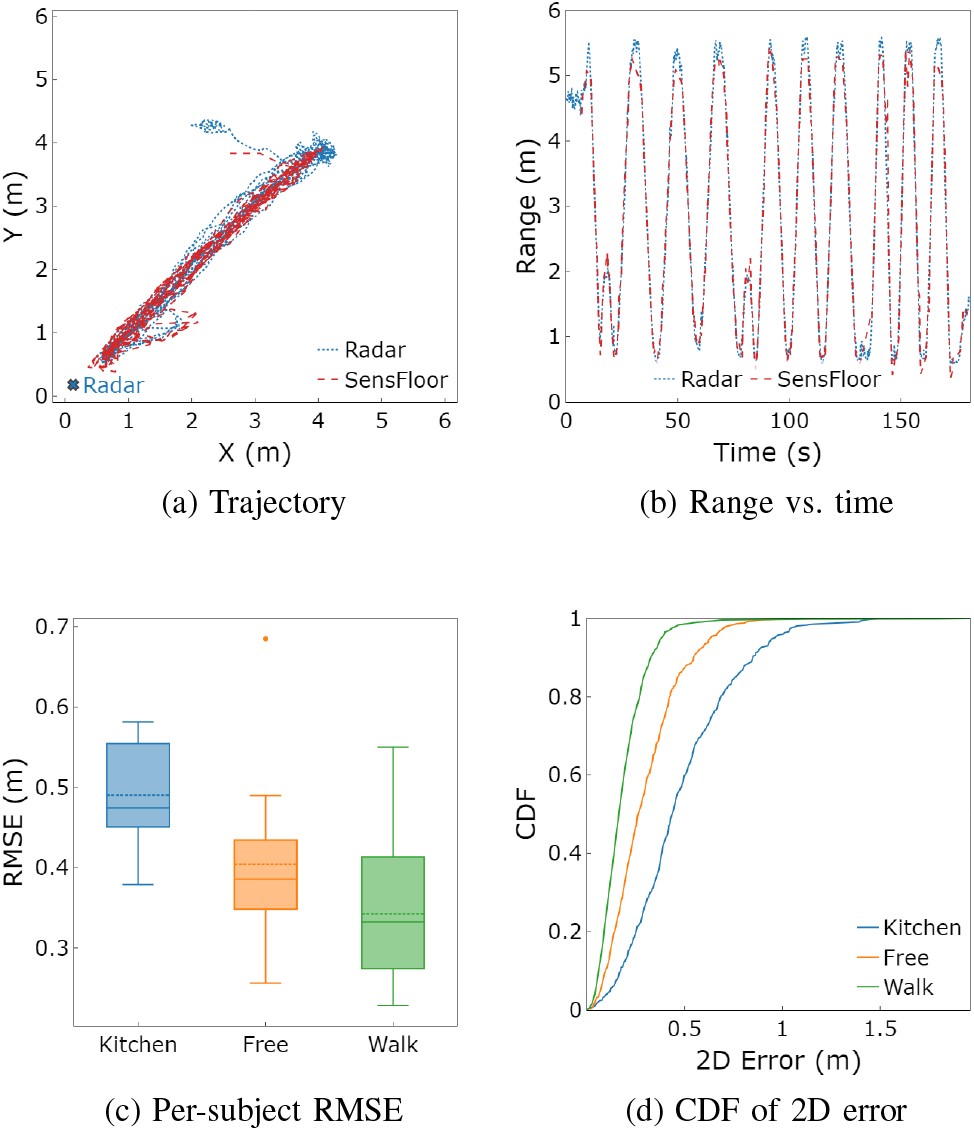
Localization results: (a) example trajectory, (b) range-time variation during structured walking, (c) per-subject RMSE by segment, (d) CDF of 2D localization error.

Across all segments and subjects, the system achieved a median 2D error of 0.24 m, with an overall mean absolute error (MAE) of 0.3 m, and root mean square error (RMSE) of 0.4 m, with 90% of errors below 0.59 m. Per-subject RMSE grouped by segment is shown in Fig. 9c. Median RMSE was highest during the kitchen task (0.47 m), followed by free exploration (0.38 m) and structured walking (0.33 m). Structured walking showed the greatest inter-subject variability, with per-subject RMSE spanning 0.23 to 0.55 m (inter-quartile range (IQR) = 0.14 m). One subject during free exploration condition produced an outlying RMSE of 0.69 m.

The cumulative distribution function (CDF) of 2D error (Fig. 9d) reflects the same segment ordering. During structured walking, 75% of errors fell below 0.31 m and 90% below 0.47 m. Free exploration achieved 75th and 90th percentile errors of approximately 0.41 m and 0.62 m. The kitchen segment exhibited the heaviest tail, with errors extending beyond 1.5 m in some cases.

The proposed system is also capable to run in real time on a Raspberry Pi 5, achieving a mean CPU usage of 26%, confirming suitability for continuous 24/7 operation on low-cost embedded hardware.

## V. Discussion

Most existing radar-based research addresses a specific sensing task, such as vital sign monitoring, activity recognition, or gait analysis. The proposed system integrates activity classification and tracking into a unified pipeline that first identifies the target’s current state and then directs processing accordingly. This offers a key practical advantage for continuous home monitoring, as only task-relevant processing is invoked at any given time. It reduces computational cost and avoids the practical challenge of retaining large volumes of raw data in real deployment. The framework addresses the infrastructure, compliance, and privacy constraints of home-based dementia monitoring, and provides a practical foundation for the per-state performance analysis discussed below.

### 1) Absence

absence detection demonstrated perfect event-level metrics, supporting further Minder deployment. An inherent onset delay of approximately 25 s is present: mean observed delays were 25.8 s for main door (besides the bed) exits and 23.7 s for toilet door exits, with toilet door exits showing substantially higher variability (SD = 24.0 s, range 5.7-96.0 s), as reported per-event in Fig. 10. In this work, delays were corrected during processing and therefore do not affect the reported metrics. However, in a real-time deployment, they would introduce a lag between the subject leaving and the system reporting an absence event. This is an inherent design trade-off, as absence confirmation requires signal variance to remain below the detection threshold for a sustained period to distinguish absence and stationary phases. Consequently, residual signal fluctuations near the target’s last known position can lead to longer delay, but the estimate would eventually converges to the correct state (e.g., the toilet door scenario for sub 01 in Fig.10). Replacing the range-dependent threshold with a 2D location-dependent threshold, concentrating the absence check within a spatially defined zone, could reduce both delay and false detection rate, and would additionally remain valid under multi-occupancy conditions.

**Fig. 10.**
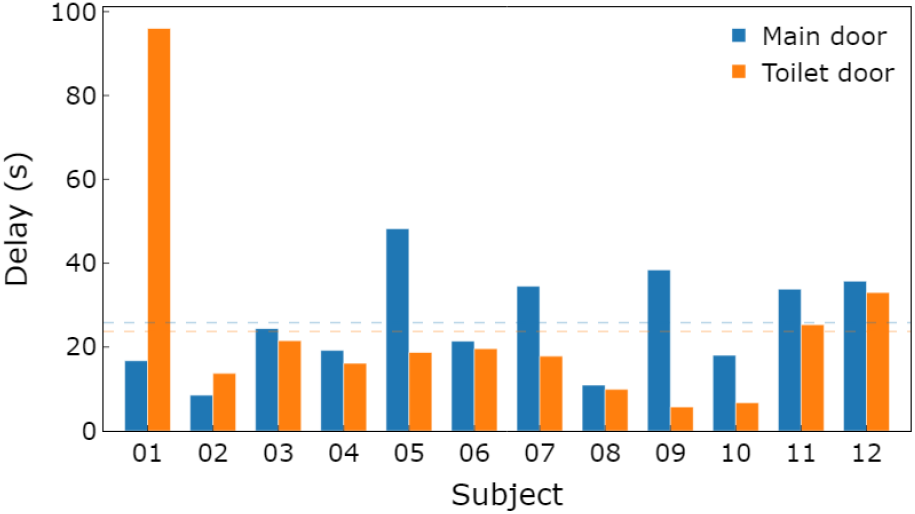
Per-subject absence confirmation delay.

### 2) Stationary Periods

the system maintained correct stationary classification throughout extended low-motion periods, including prolonged lying on the bed. Three methodological points contextualize the perfect frame-level metrics. First, stationary periods constitute the longest-duration class (167,452 of 230,045 frames), so isolated misclassifications had limited quantitative impact. Second, labeling was conservative: only periods exceeding 25 s and confirmed against camera annotation were included. Third, fine hand movements are suppressed by the torso velocity gate, particularly at greater range or when the target faces away from the radar, and are therefore excluded from the stationary definition. To capture absolute stillness required for clinically accurate vital sign monitoring, a more sensitive confirmation method would be needed, such as the confidence-level-based confirmation proposed in [23].

### 3) Walking

walking detections achieved a duration-weighted sensitivity of 0.914 ±0.026, confirming that missed events are short marginal bouts rather than clinically significant ambulation. For long-term 24/7 monitoring, a conservative detection strategy is preferable, as tolerating a higher false negative rate yields a cleaner walking dataset. Unlike spatial trajectories that map broad behavioral patterns, simple walking event counts offer less clinical value than the detailed gait and identity metrics extracted from reliable, high-confidence walking periods. Additional false negatives arose from cross-range walking and table-obstacle movements, which produced degraded foot trajectory. Per-segment sensitivity (Fig. 11) shows structured walking achieved the highest value, while kitchen task performance was degraded by short-bout dominance, directly characterizing expected variation across real-home activity contexts. Two false positive sources are worth noting: (a) posture changes during lying phase generated radial motion and foot trajectory values that confused the detector; (b) during kitchen tasks and free exploration, fast hand motions following a true walking segment, such as reaching for upper cabinets, produced spurious foot trajectory and radial motion.

**Fig. 11.**
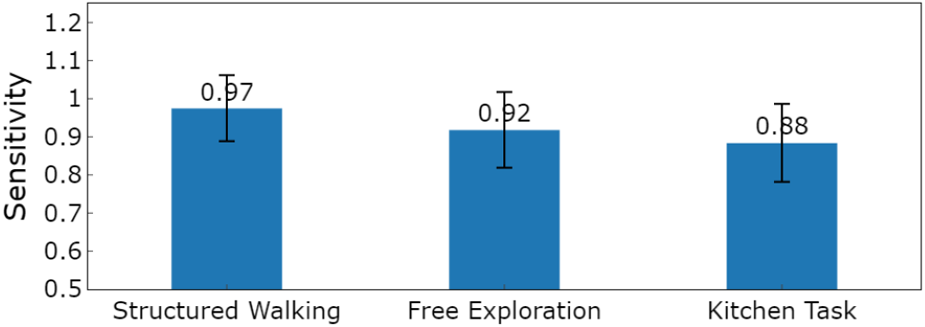
Walking detection sensitivity across activity segments.

### 4) Localization

localization errors arose from three main sources: (a) SensFloor grid resolution of 0.5 m imposes a quantization floor on achievable accuracy; (b) foot pressure versus torso position measurements introduce a systematic offset varying with posture and gait; (c) SensFloor provides no output during stationary phases, explaining elevated errors during the kitchen task where subjects frequently stood still yet with torso movements. Accuracy was further affected by the radar’s ±45^*°*^ azimuth limit, placing the toilet and bedside areas near the detection boundary, where angular estimation error increases. This is consistent with the outlier observed in Fig. 9c, where subject 12 spent a noticeable amount of time exploring the edge areas.

The reported RMSE of 0.40 m is consistent with comparable single-radar mmWave systems (0.20-0.40 m under controlled conditions [24] [25]), and compares favorably with wearable-based systems (mean 1.66 m [26]), while avoiding the compliance burden relevant to dementia monitoring. For the room-level and zone-level localization tasks required in dementia care, the achieved accuracy is considered sufficient. Stationary-phase localization remains the primary uncharacterized deployment risk: torso posture changes during lying and sleep produce variable radar returns, and the largest return may not originate from the expected body center, as SensFloor provides no ground truth outside active walking.

### 5) Limitations and Future Work

validation involved a small cohort of healthy young participants, therefore does not fully capture performance across diverse body types, mobility patterns, and extended time frames. Additional gaps remain before deployment: overnight and sleep-phase stationary behavior has not been validated, which is critical for 24-hour Minder deployment, and multi-occupancy scenarios remain unaddressed. Future work will focus on an extended pilot in real Minder homes with wearable ground truth over continuous monitoring periods, addressing temporal stability, overnight behavior, and naturalistic gait simultaneously.

## VI. Conclusion

Bridging the gap between laboratory sensing and scalable 24/7 home healthcare, this work presents a fully autonomous radar framework capable of extracting continuous, clinically relevant behavioral patterns from unconstrained home environments. The proposed system performs real-time activity classification (stationary, walking, absent) alongside target localization, and is explicitly designed for continuous 24/7 operation without manual intervention. It addresses key real-world deployment challenges: generalizable thresholding across subjects and environments, robust absence detection for stationary targets, and reliable walking detection during naturalistic daily activities. Validation demonstrated 0.98 frame-level accuracy, event-level F1 *>* 0.93 across all states, and 0.40 m localization RMSE during movement. False positive sources were characterized and attributed to identifiable failure modes, supporting confidence in deployment reliability under naturalistic conditions. Results demonstrate readiness for transition to long-term real-world deployment within the Minder platform. Overnight validation, multi-occupancy handling, and naturalistic elderly gait characterization form the concrete roadmap for next stage of clinical evaluation.

## Acknowledgements

This work is supported by the UK Dementia Research Institute [award number UK DRI-7204] through UK DRI Ltd, principally funded by the Medical Research Council, and additional funding partner Alzheimer’s Society

## References

[1] D. Carter. (2016) Fix dementia care: Homecare. Alzheimer’s Society. Policy Officer, Alzheimer’s Society.

[2] Alzheimer’s Society. (2023) Two-thirds of gps want to prescribe tech for patients diagnosed with dementia.

[3] A. Mannion. (2024) Carers say tech is crucial for people with dementia but they need digital skills support. AbilityNet.

[4] T. G. Stavropoulos, A. Papastergiou, L. Mpaltadoros, S. Nikolopoulos, and I. Kompatsiaris, “Iot wearable sensors and devices in elderly care: A literature review,” Sensors, vol. 20, no. 10, p. 2826, 2020.

[5] S. Kekade et al., “The usefulness and actual use of wearable devices among the elderly population,” Computer Methods and Programs in Biomedicine, vol. 153, pp. 137–159, 2018.

[6] Y. Cai, B. Guo, F. Salim, and Z. Hong, “Towards generalizable human activity recognition: A survey,” arXiv preprint arXiv:2508.12213, 2025.

[7] A. K. Alhazmi, M. A. Alanazi, A. H. Alshehry, S. M. Alshahry, J. Jaszek, C. Djukic, A. Brown, K. Jackson, and V. P. Chodavarapu, “Intelligent millimeter-wave system for human activity monitoring for telemedicine,” Sensors, vol. 24, no. 1, 2024.

[8] Y. Liu, J. Zhang, Y. Chen, W. Wang, S. Yang, X. Na, Y. Sun, and Y. He, “Real-time continuous activity recognition with a commercial mmwave radar,” IEEE Transactions on Mobile Computing, vol. 24, no. 3, pp. 1684–1698, 2024.

[9] C.-Y. Hsu, “Passive sensing of user behavior and well-being at home,” Thesis, 2020.

[10] S. M. M. Islam, “Radar-based remote physiological sensing: Progress, challenges, and opportunities,” Frontiers in Physiology, vol. 13, p. 955208, 2022.

[11] UK Dementia Research Institute, “Minder — uk dri care research & technology centre,” https://www.ukdri.ac.uk/centres/care-research-technology/minder, accessed: 2026-01-26.

[12] A. Soumya, C. Krishna Mohan, and L. R. Cenkeramaddi, “Recent advances in mmwave-radar-based sensing, its applications, and machine learning techniques: A review,” Sensors, vol. 23, no. 21, 2023.

[13] S. M. Kahya, M. S. Yavuz, and E. Steinbach, “Hood: Real-time human presence and out-of-distribution detection using fmcw radar,” IEEE Transactions on Radar Systems, vol. 3, pp. 44–56, 2024.

[14] A. Ninos, J. Hasch, M. Heizmann, and T. Zwick, “Radar-based robust people tracking and consumer applications,” IEEE Sensors Journal, vol. 22, no. 4, pp. 3726–3735, 2022.

[15] H. Yang, D. Zhang, X. Zhang, J. Xiong, Z. Fan, W. Ning, W. Chen, F. Zhang, Z. Han, and D. Zhang, “From spatial domain to temporal domain: Unleashing the capability of cfar for mmwave point cloud generation,” Proceedings of the ACM on Interactive, Mobile, Wearable and Ubiquitous Technologies, vol. 9, no. 2, pp. 1–29, 2025.

[16] C. Hadjipanayi, M. Yin, A. Bannon, Z. Chen, and T. G. Constandinou, “Towards radar-agnostic gait analysis across uwb and fmcw systems,” arXiv preprint arXiv:2601.04415, 2026.

[17] C. Hadjipanayi, M. Yin, A. Bannon, A. Rapeaux, M. Banger, S. Haar, T. S. Lande, A. H. McGregor, and T. G. Constandinou, “Remote gait analysis using ultra-wideband radar technology based on joint range-doppler-time representation,” IEEE Transactions on Biomedical Engineering, vol. 71, no. 10, pp. 2854–2865, 2024.

[18] M. A. Richards, J. A. Scheer, and W. A. Holm, Principles of modern radar: basic principles. IET, 2010.

[19] Y. Bar-Shalom, X. R. Li, and T. Kirubarajan, Estimation with applications to tracking and navigation: theory algorithms and software. John Wiley & Sons, 2001.

[20] P. Nallabolu, L. Zhang, H. Hong, and C. Li, “Human presence sensing and gesture recognition for smart home applications with moving and stationary clutter suppression using a 60-ghz digital beamforming fmcw radar,” Ieee Access, vol. 9, pp. 72 857–72 866, 2021.

[21] J. Perry and J. Burnfield, Gait analysis: normal and pathological function. CRC Press, 2024.

[22] M. Crook-Rumsey, S. J. Daniels, S. Abulikemu, H. Lai, A. Rapeaux, C. Hadjipanayi, E. Soreq, L. M. Li, J. Bashford, J. Jeyasingh-Jacob et al., “Multicohort cross-sectional study of cognitive and behavioural digital biomarkers in neurodegeneration: the living lab study protocol,” BMJ open, vol. 13, no. 8, p. e072094, 2023.

[23] Z. Chen, A. Bannon, A. Rapeaux, and T. G. Constandinou, “Towards robust, unobtrusive sensing of respiration using ultra-wideband impulse radar for the care of people living with dementia,” bioRxiv, pp. 2020–12, 2020.

[24] Z. Xing, P. Chen, J. Wang, Y. Bai, J. Song, and L. Tian, “Millimeter-wave radar detection and localization of a human in indoor complex environments,” Remote Sensing, vol. 16, no. 14, 2024.

[25] A. Venon, Y. Dupuis, P. Vasseur, and P. Merriaux, “Millimeter wave fmcw radars for perception, recognition and localization in automotive applications: A survey,” IEEE Transactions on Intelligent Vehicles, vol. 7, no. 3, pp. 533–555, 2022.

[26] K. Roohi and A. R. Fekr, “A comparative analysis of indoor localization technologies,” Computer Networks, vol. 270, p. 111527, 2025.

